# Influence of hyperparameters on the performance of deep learning-based microrobotic localization under phantom tissue

**DOI:** 10.1101/2025.08.13.670026

**Authors:** Johanna Hoppe, Richard Nauber, David Castellanos Robles, Jürgen Czarske, Mariana Medina-Sánchez

**Affiliations:** Micro- and Nano-Biomedical Engineering Group, Leibniz Institute for Solid State and Materials Research Dresden, Institute for Emerging Electronic Technologies, Dresden, Germany; Micro- and NanoBiosystems, Center for Molecular Bioengineering (B CUBE), Dresden University of Technology, 01062, Dresden, Germany; Ikerbasque, CIC NanoGUNE, Donostia-San Sebastian, Spain; Laboratory of Measurement and Sensor System Technique (MST), TU Dresden, Helmholtzstrasse 18, 01069 Dresden, Germany

## Abstract

For the effective operation of medical microrobots within living organisms and precise targeting, it is imperative to employ imaging techniques closely integrated with real-time deep-tissue tracking methods. However, due to a typically low Signal-to-Noise ratio images with strong background, it is hard for traditional tracking methods to achieve sufficient accuracy. This challenge can be addressed by deep learning-based tracking with a real-time detection model. However, a multitude of design choices and Hyperparameters influence the performance. In this study we compared the influence of the hyperparameters and model architecture versions of the “you only look once” (YOLO) network. We use experimental data from a magnetic microrobot imaged with Photoacoustics through 5 mm phantom tissue to evaluate the tracking in comparison with an optical reference. The deep-learning based methods consistently achieved lower missing-detection ratios. Regarding the Root Mean Square localization error, we observed that increasing the weight of the box loss function and utilizing the distribution focal loss can enhance the performance by 10%. Furthermore, it can be seen that YOLOv9 consistently outperformed its predecessor YOLOv8. This study quantifies the robustness of deep-learning based tracking of medical microrobots under tissues.

## I. Introduction

The field of medical microrobots is undergoing rapid evolution with the perspective of reducing the invasiveness of medical interventions in the future [1][2][3][4]. For operating microrobots in vivo, it is of paramount importance to be able to detect and track them under deep tissue in real-time. Photoacoustic imaging is capable of meeting that requirement without ionizing radiation [5][6][3], however a challenge of automatic tracking based on the very low signal-to-noise ratio (SNR) images remains.

An object detection neural network could, in a supervised learning process, be trained to differentiate between background noise and the photoacoustic signal of the actual magnetic microrobot. The CLC system (Closed-Loop-Control system) is responsible for the cooperation of every component controlling the movement of the magnetic micro-robot, and thus depends on the reliable real-time capability of the tracking and detection of the microrobot. The objective of this work is to reveal the influence of model architecture and loss function on the detection performance through a systematic study [7].

## II. Methods

### A. Deep Learning-based Object Detection Models

The latest version of the YOLO algorithm, version 9, was introduced in February 2024. It exhibits enhanced efficiency, accuracy, and adaptability in comparison to its predecessor, version 8. This enables promise more robust detection and more efficient learning. The functional enhancements are supported by a novel model structure that integrates a PGI (programmable gradient information) and GELAN (Generalized Efficient Layer Aggregation Network) [8].

A more detailed examination of the differences between the YOLOv8 and YOLOv9 models is presented in Figure 1. In this study, we utilized both the YOLOv8-s and YOLOv9-e models. The overall complexity of YOLOv9-e is greater than that of YOLOv8, with an increased number of parameters (58M vs. 11.1M) and floating point operations (FLOPs; 192.5 vs. 28.6). The input image size is identical in both models, with dimensions of 640 by 640 pixels. Both models share a similar base structure, comprising a descending backbone, a feature-extracting neck, and a decoupled head for classification and bounding box prediction [9].

**Fig. 1:**
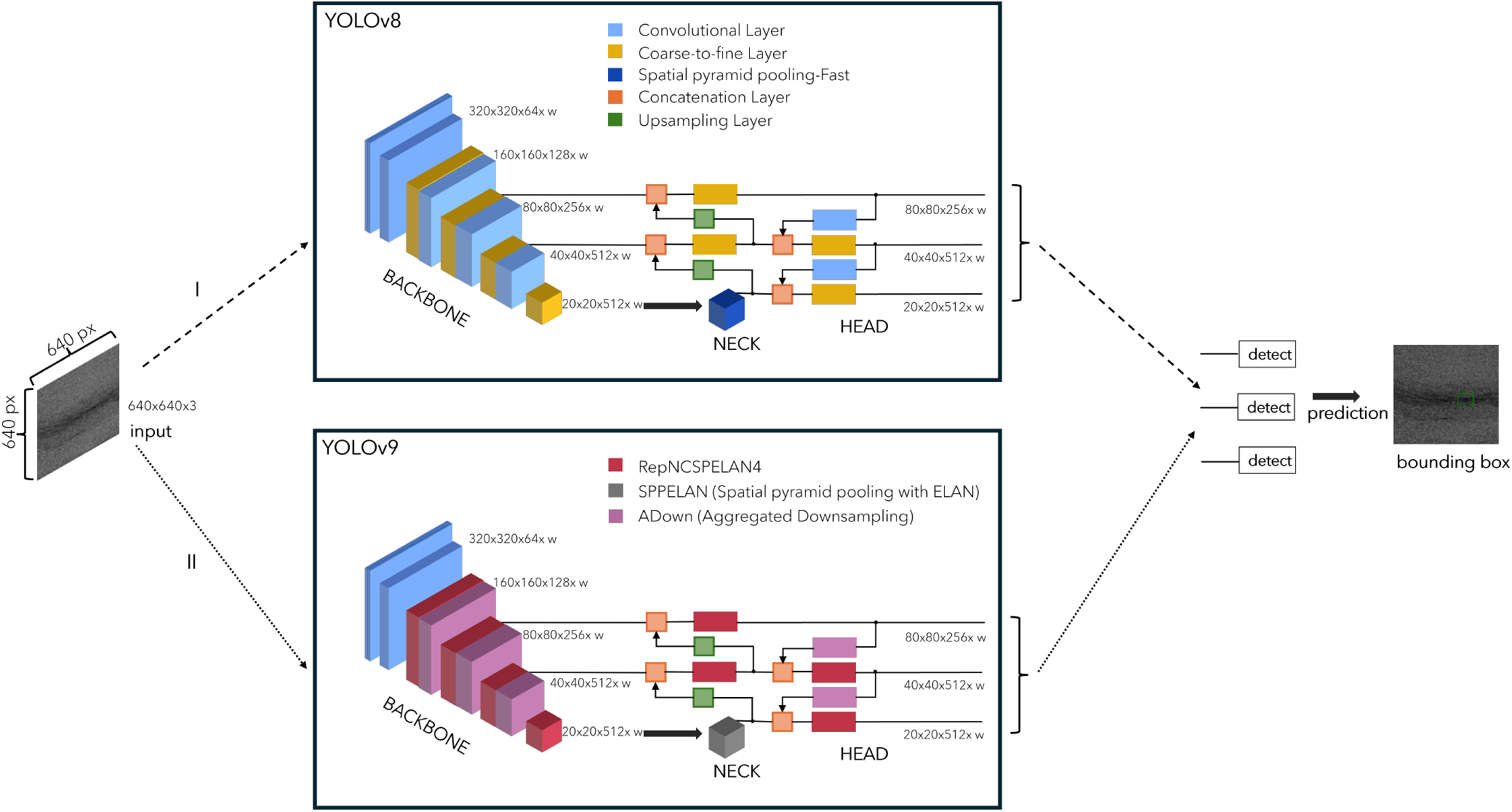
YOLOv8 and YOLOv9 model structure based on [10] and [11]

**Fig. 2:**
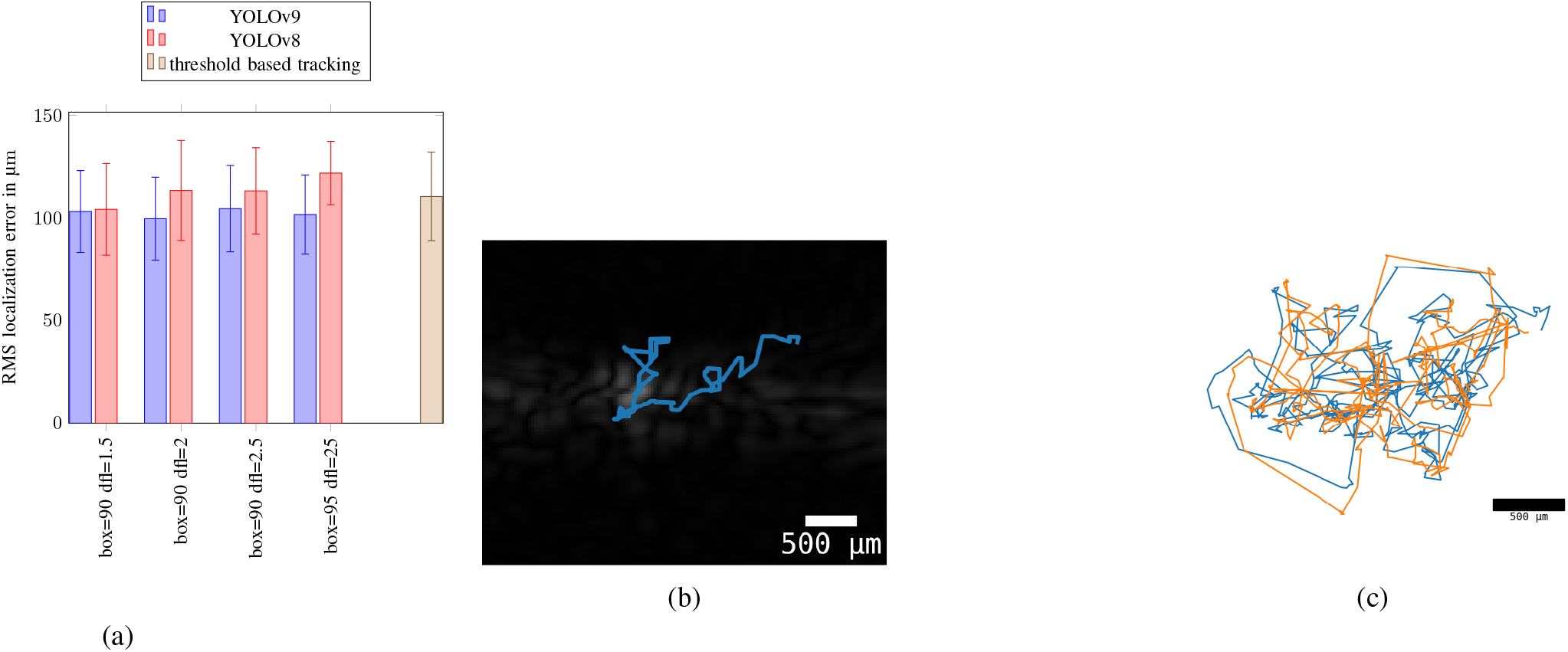
a) RMS localization error for different hyperparameter sets; b) tracked trajectory in live experiment for box=90 and dfl=20 with YOLOv9; c) trajectory comparison of optical tracking and tracking with model from b

The main differences between the two models are presented in Figure 1. These differences are present in the backbone and head. In YOLOv8, the C2f (coarse to fine) layers were exchanged for further two way feature extraction (RepNCSPELAN4) layers in YOLOv9. With the exception of the initial two convolutional layers, the remaining convolutional layers were replaced with ADown layers for downsampling. In the neck, the conventional SPPF block in v8 was supplanted by the enhanced SPP (Spatial pyramid pooling) structure with ELAN. This new structure is capable of capturing multi-scale contextual information [9].

### B. Loss Function types

In order to achieve optimal results, it is essential that the loss function used to train the model is matched to the specific detection task. YOLO models are designed for multiclass detection tasks. In our case, we are attempting to detect a single class with different shapes. The YOLO model incorporates three distinct loss functions, with the recognition of the class being performed by BCE (binary cross entropy) in order to reconcile the classification loss. The regression branch, which attempts to regress the bounding box, comprises the distribution focal loss (DFL) and the intersection over union (IoU) loss, also known as boxloss. DFL is beneficial in the context of imbalanced datasets and bounding box regression, particularly when the data provided to the model exhibits unclear boundaries. IoU represents a metric for the accuracy of the bounding box placement. The weighted sum of the three functions is the final loss [12].

The objective is to place the bounding box around the center of the object and minimize its size simultaneously. This is achieved by regressing the bounding box with adjustments to the weights of the regression branch loss functions, dfl and IoU. In this case, the influence of the cls function is minimal.

## III. Results

This work investigates the effect of loss function weights on prediction quality, measured by Root Mean Square (RMS) localization error and the number of detected frames relative to the total number of frames. The experiment is a magnetic microrobot (250 250 µm cuboid magnet covered in Alginate) moved by a magnetic field in CLC (Closed-Loop-Control) setup, through 5mm of gelatin representing a tissue-like factor causing dispersion of the signals, as described in [7]. Both models, YOLOv8 and YOLOv9, were trained under the same conditions: a dataset containing 5583 training examples, 682 validation examples and 452 test images. The dataset used for the training process is completely independent from the experiment data. A comparison was conducted between the two models, with different values for the box loss and the dfl weights, in order to examine the changes in the model accuracy and confidence. Comparing the results for both model types as shown in table 1, it is visible that YOLOv9 is performing better than YOLOv8, both in terms of accuracy, the RMS localization error, and the confidence, and missed detection ratio. the best hyperparameter configuration used with YOLOv9 is: a box loss weight of 9, a dfl weight of 2 and a cls loss weight of 0.5. Compared to the default values: box-loss 7.5, dfl-loss 1.5, and cls=0.5, confirmed our original thesis, that the choice of weight for the loss function influences the training outcome dependent on the datatype represented in the dataset. Furthermore, the results confirmed our original hypothesis that increased box loss weights and dfl weights serve the purpose of a single class detection and tracking with different representations. Our hypothesis was not only validated but also demonstrated to enhance tracking performance compared to threshold based tracking. The tracking without machine learning achieved an RMS localization error of 110.468 µm and the missing detection rate was 10 of 513, while our best model realized an RMS localization error of 99.607 µm and missed detection rate. This study quantifies the robustness of deep-learning based tracking of medical microrobots under tissue.

**TABLE I:** Comparison of the prediction results of YOLOv8 and YOLOv9 with different weights for the loss functions.

| YOLO Version | Box | DFL | CLS | RMS in $\mu\text{m}$ | $\frac{\text{Lost Items}}{\text{Total Items}}$ |
| --- | --- | --- | --- | --- | --- |
| threshold based | | | | $110.465 \pm 21.586$ | 10/515 |
| v9 | 9 | 1.5 | 0.5 | $103.116 \pm 19.951$ | 0 |
| | | 2 | | $99.607 \pm 20.210$ | 0 |
| | | 2.5 | | $104.531 \pm 21.047$ | 0 |
| v9 | 9.5 | 2.5 | 0.5 | $101.643 \pm 19.257$ | 0 |
| v8 | 9 | 1.5 | 0.5 | $104.177 \pm 22.342$ | 0 |
| | | 2 | | $113.375 \pm 24.361$ | 0 |
| | | 2.5 | | $113.171 \pm 21.023$ | 0 |
| v8 | 9.5 | 2.5 | 0.5 | $121.836 \pm 15.447$ | 1/515 |

## ACKNOWLEDGMENT

The authors like to thank David Weik and Zehua Dou from Laboratory of Measurement and Sensor System Technique (MST), TU Dresden, for creative and productive discussions.

